# STAMP-Based Digital CRISPR-Cas13a (STAMP-dCRISPR) for Amplification-Free Quantification of HIV-1 Plasma Viral Load

**DOI:** 10.1101/2022.10.13.512138

**Authors:** Reza Nouri, Yuqian Jiang, Anthony J. Politza, Tianyi Liu, Wallace Greene, Jonathan Nunez, Xiaojun Lance Lian, Weihua Guan

**Affiliations:** Department of Electrical Engineering, Pennsylvania State University, University Park, Pennsylvania 16802, United States; Department of Biomedical Engineering, Pennsylvania State University, University Park, Pennsylvania 16802, United States; Department of Pathology, Penn State College of Medicine, Hershey, PA 17033, United States; Department of Medicine, Penn State College of Medicine and Milton S. Hershey Medical Center, Hershey, Pennsylvania 17033, United States; Huck Institutes of the Life Sciences, Pennsylvania State University, University Park, Pennsylvania 16802, United States; Department of Biology, Pennsylvania State University, University Park, Pennsylvania 16802, United States

## Abstract

The development of new nucleic acid techniques to quantify HIV RNA in plasma is critical for identifying the disease progression and monitoring the effectiveness of antiretroviral therapy. While RT-qPCR has been the gold standard for HIV viral load quantification, digital assays could provide an alternative calibration-free absolute quantification method. Here, we report the development of a <u>s</u>elf-digitalization <u>t</u>hrough <u>a</u>utomated <u>m</u>embrane-based <u>p</u>artitioning (STAMP) technique to digitalize the CRISPR-Cas13 assay (dCRISPR) for amplification-free and absolute quantification of HIV-1 viral RNAs. The analytical performances of STAMP-dCRISPR were evaluated with synthetic HIV-1 RNA, and it was found samples spanning 4 orders of dynamic range between 100 aM to 1 pM can be quantified as fast as 30 min. We also examined the overall assay from RNA extraction to STAMP-dCRISPR quantification with spiked plasma samples. The overall assay showed a resolution of 42 aM at a 90% confidence level. Finally, a total of 20 clinical plasma samples from patients were evaluated with STAMP-dCRISPR. The obtained results agreed well with the RT-qPCR. Our result demonstrates a new type of easy-to-use, scalable, and highly specific digital platform that would offer a simple and accessible platform for amplification-free quantification of viral RNAs, which could be exploited for the quantitative determination of viral load for an array of infectious diseases.

---

Acquired immunodeficiency syndrome (AIDS) caused by human immunodeficiency virus (HIV) infection, a notorious fatal epidemic, has led to millions of deaths worldwide since its origin^1^. Although AIDS-related annual mortality has reduced by 33% in the past decade due to the application of antiretroviral therapies and advanced HIV diagnosis, the number of new HIV infections remains high (for instance, 1.5 million in 2020 globally), which is estimated to cost billions of dollars for AIDS therapy ^2^. Since AIDS patients at early stages tend to present no obvious symptoms but can still be infectious, early awareness of infection enables timely treatment for exposed patients and prevents further transmission ^3^. Viral load quantification of the HIV-1 RNA not only identifies the progression of the disease in a patient but also could be employed to monitor the trends in large populations of patients ^4-6^. Therefore, nucleic acid tests (NAT) that have been utilized for viral load quantification hold tremendous promise in AIDS diagnosis ^7^. One of the major techniques for viral load quantification of HIV is the reverse transcriptase quantitative polymerase chain reaction (RT-qPCR) due to its accessibility and high sensitivity ^4,8^. Although RT-qPCR has been the gold standard for detecting the HIV-1 RNA ^9,10^, the emerging clustered regularly interspaced short palindromic repeats (CRISPR) based technology has taken immense attention for nucleic acid tests due to its high sensitivity and specificity ^11,12^.

Since the discovery of Cas9 proteins for gene editing, the CRISPR technology has taken center stage in biotechnology ^13^. Recently, the discovery of the collateral cleavage in other Cas proteins like Cas12 ^14^ and Cas13 ^15^ made it possible to translate the sequence-specific targeting to other detectable signals, which has led to the increasing emergence of CRISPR-mediated biosensors ^15-28^. Among these CRISPR-mediated assays, a preamplification step is often required to boost the limit of detection and time to results performance ^23,29^. However, preamplification complicates the assay setup, increases the assay time, raises the risk of contamination, and could introduce false-negative or -positive results due to amplification errors ^30^. We previously investigated different strategies to boost the CRISPR assay performance ^31^, such as the use of Cas proteins with higher cleavage activity ^32^, the use of multiple crRNA in the reaction ^33,34^, the use of a sensitive readout system ^35^, and the reaction digitalization ^34,36-47^. Among these techniques, we found only the digitalization method could match the limit of detection (attomolar range) and the fast turnaround time (less than 1 hour) of preamplification-coupled CRISPR assays ^31^.

So far, various digitalization techniques have been introduced. For instance, water in oil droplets generated by T-junction ^48^, flow focusing ^49,50^, and centrifugation ^51^ have been used for digitalization. Furthermore, digital assays have been performed inside numerous microchambers fabricated by polydimethylsiloxane (PDMS) or glass chambers. Partitioning of the assay inside these chambers has been achieved using vacuum ^52,53^, pressure ^54^, SlipChip ^55^, hydrophilic patterns ^56,57^, or self-digitization ^58^. While these techniques have been optimized and improved considerably, complicated fluidic control systems and complex micro and nanofabrication processes are required for them. Therefore, developing a platform to eliminate complicated fluidic control and fabrication processes would be desirable for highly accessible digital assay systems.

In this study, we demonstrated a self-digitalization method through an automated membrane-based partitioning (STAMP) and developed a STAMP-based digital CRISPR-Cas13a (STAMP-dCRISPR) for the absolute quantification of HIV-1 viral load. We first established a stamping technique to digitalize the Cas13a assay inside a commercial track-etched polycarbonate (PCTE) membrane without a complicated fluidic control process (*e.g*., pump, vacuum, and valve). To optimize the Cas13a assay, we studied the effect of different CRISPR RNA (crRNA) design and assay reaction time on the sensing performance. The absolute quantitative performance, the limit of detection and the dynamic range were evaluated by quantifying the synthetic HIV-1 RNA at different concentrations. We also examined the overall assay from plasma RNA extraction to STAMP-dCRISPR with spiked HIV-1 plasma samples. Finally, the clinical applicability of the STAMP-dCRISPR was demonstrated in the absolute quantification of 20 HIV-1 patient samples.

## Results

### STAMP device and characterization

In order to achieve self-digitalization without complicated fluidic control, we developed the STAMP method to digitalize the assay. In this method, a commercial polycarbonate track-etched (PCTE) membrane was utilized for digitalization. This type of commercial membrane consists of a high density of micro/nanopores with uniform pore sizes ranging from 10 nm to 30 μm [29]. **Figure 1a** illustrates a top and side view of the assembled stamp device where the membrane is sandwiched between a polymethyl methacrylate (PMMA) holder and a thin tape (70 μm thickness). In addition, **Figure 1a** depicts an image of the transparent and flexible PCTE membrane with a diameter of 1.3 cm. We characterized the pore size distribution by examining 5 different membranes. As shown in **Figure 1b**, the average pore size was measured as 24.6±1.6 μm, and the pore density was determined to be 9895±531 pores/cm^2^.

**Figure 1.**
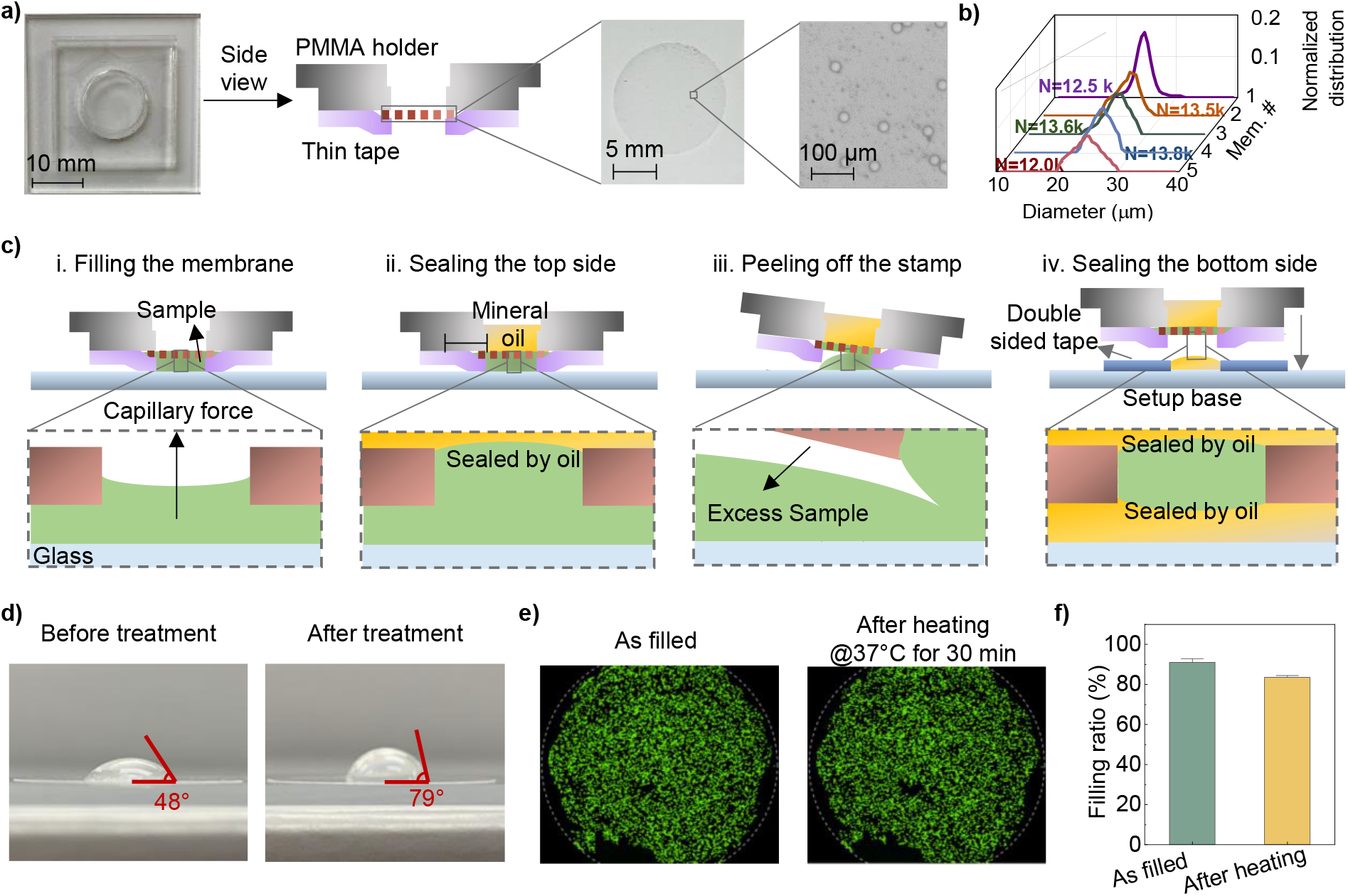
STAMP device characterization and filling process. a) Different components of the STAMP system along with a top-side view of the assembled device and images of the commercial PCTE membranes. b) Pore size distribution of five different membranes and their total number of pores. d) STAMP process. **i**. The process starts by placing the stamp device on top of the sample. **ii**. The top side of the system is sealed by adding mineral oil. **iii**. The stamp is removed from the glass to eliminate the excess liquid from the bottom of the membrane. **iv**. The stamp is placed on the setup base (consisting of glass, double-sided tape, and mineral oil) to seal the bottom side of the system. d) Chemical treatment to remove the polyvinylpyrrolidone (PVP) coating from the PCTE membrane. The contact angle of a water droplet on top of the membrane increased from 48 to 79 degrees after treatment, confirming the effectiveness of the PVP removal process. e) Fluorescent images of a membrane demonstrating the filling of the membrane using STAMP before and after 30 minutes of heating at 37 °C. All filled pores are labeled with a filled green circle to demonstrate the filling process. f) Measured filling ratio of the membranes before and after 30 minutes of heating at 37 °C. We used a bright image of the membrane to estimate the total number of pores.

The operation of the stamp device only requires 4 simple manual steps. In the first step (**Figure 1c-i**), the analyte sample droplet was deposited on top of a glass surface, and the stamp was slowly placed on top of it. To ensure the filling process, only 8 μL of the sample was required, which is 33% more than the spacing volume of 6 μL between the membrane and glass surface. Once in contact, the surface tension between the sample and pore walls causes a capillary action that forces the sample into the membrane’s pores. After 60 seconds of soaking, 60 μL of mineral oil was added to the top chamber to seal the top surface of the membrane (**Figure 1c-ii**). An inspection of the stamp confirmed that all pores were successfully filled despite that there were excessive liquids underneath the membrane (**Figure S1a**). To remove these excessive samples, one only needs to peel off the stamp from the glass surface (**Figure 1c-iii**). The as-purchased membranes were coated with polyvinylpyrrolidone (PVP) which renders the surface hydrophilic. To facilitate the excessive liquid removal, this hydrophilic coating was removed by dipping the membranes in 10% acetic acid for 30 minutes and heating at 140 °C for 60 minutes in a vacuum oven. **Figure 1d** shows that the contact angle increased from 48 to 79 degrees after this chemical treatment, confirming the PVP removal process. Since the glass surface is hydrophilic and the PCTE membrane surface is hydrophobic, the excess liquid would remain on the glass and be removed from the membrane surface. In the pore areas, the surface tension overcomes the liquid intermolecular forces and holds the sample inside the pores. An examination of the stamp confirms this process for effectively removing the excess liquid while maintaining the digitalized samples (**Figure S1b**). Lastly, the stamp was placed on top of a customized base with prefilled mineral oil (**Figure 1c-iv**) to form a fully sealed digital system for further reaction.

To evaluate the membrane filling process and evaporation under heating procedures, we measured the filling ratio (total number of filled pores per total number of pores) of the final sealed membrane before and after 30 minutes of heating at 37 °C. **Figure 1e** and **Figure 1f** illustrate representative fluorescent images of the membrane and the measured filling ratio before and after the heating procedure. The average filling ratio before the heating was measured as 91.09%. Usually, some parts of the membranes do not fill, which could be caused by the sample intermolecular forces overcoming the surface tension when the stamp is removed from the glass. After 30 minutes of heating at 37 °C, we observed evaporations in some parts of the membrane where the filling ratio reduced to 83.54%. Those unfilled pores show no fluorescence signals at all and can be easily distinguished from the pores with negative reactions (which exhibit weak fluorescence signals, **Figure S2**). To improve the accuracy of the absolute quantification, we only considered the filled pores as the total number of reactions in our system.

### Automated HIV-1 STAMP-dCRISPR system and its noise floor

After the development of the STAMP device, we set out to develop a platform to utilize the STAMP for running the digital CRISPR (dCRISPR) assay for HIV-1 viral load quantification. The Cas13a reaction mix was digitalized inside the membrane using the STAMP technique (**Figure 2**a). With binding to the specific RNA-guided target, Cas13a proteins become activated and perform trans-cleavage on the surrounding FQ-labeled single-stranded reporter ^14^ (**Figure 2b)**.

**Figure 2.**
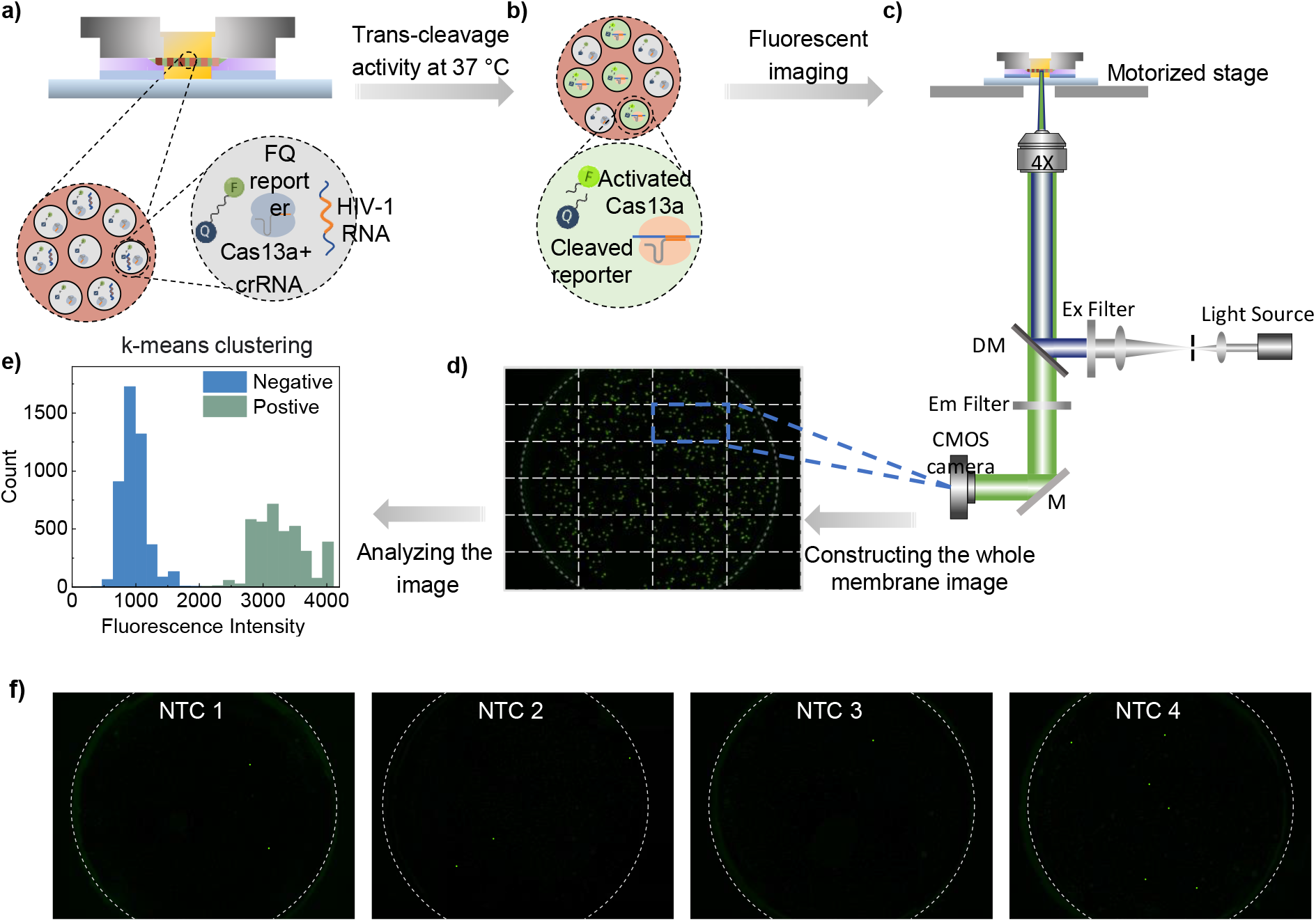
Utilization of STAMP device for running the digital CRISPR assay for HIV-1 viral load quantification. a) Digitalization of CRISPR-Cas13a assay including HIV-1 RNA, Cas13a and crRNA complex, fluorophore quencher (FQ)-labeled single-stranded RNA reporters. b) Trans-cleavage activity of the activated Cas13a proteins (after binding with HIV-1 RNAs) on non-target surrounding FQ RNA reporters. Cleavage of the reporters results in the FAM fluorescence illumination. c) Fluorescent imaging setup. d) The fluorescent image of a whole membrane stitched from 24 images taken by the microscope. e) Clustering the positive and negative pores based on their fluorescent intensity using a k-means clustering algorithm. f) Fluorescent images illustrating positive and negative pores at 4 negative control cases. All positive pores are labeled with a filled green circle for better demonstration.

Fluorescence images of the membranes were taken by a fluorescence microscope with a motorized stage to cover the whole membrane (**Figure 2c**). The light source wavelength was filtered to 480 nm using an excitation filter and redirected to the sample using a dichroic mirror. Afterward, the emitted light from the sample was obtained by CMOS camera after filtration. 24 images were taken and stitched together to cover the whole membrane area (**Figure 2d**). The acquired images were analyzed to distinguish positive from negative pores based on the fluorescent intensity emitted from each pore. We utilized a *k*-means clustering algorithm to differentiate between positive and negative pores ^59^ (**Figure 2e, Figure S3**). The Poisson statistics was utilized to quantify the number of HIV-1 RNA targets without external references. With *n* total number of reactions, the positive pore ratio (*PPR*) is defined as *PPR=m/n*, where *m* is the number of positive reactions. Based on the Poisson statistics, the concentration of the sample could be estimated as:

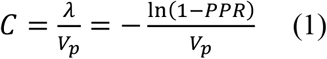

where *λ* is the expected number of targets in each pore, and *V*_*p*_ is the average volume of the pores.

We examined the no-target control (NTC) cases to obtain the background noise of STAMP-dCRISPR. **Figure 2f** presents the fluorescent images of the 4 NTC cases. While no targets were added in these cases, few positive pores were detected (**Table S1**). Multiple factors could cause the background noise in our systems, such as non-specific reporter cleavage ^60,61^, imaging ^62,63^, and post-processing inaccuracy ^62^. The system background noise (defined as μ_NTC_+3σ_NTC_) was measured as 0.00093, where μ_NTC_ and σ_NTC_ are the averages and standard deviation of the *PPR* in the negative cases measured.

### Design and optimization of HIV-1 Cas13 assay

To optimize the Cas13 crRNA design, we initially designed five crRNAs along the HIV-1 genome (red rectangles in **Figure 3a, Table S2**). In addition, we synthesized five 100 nucleotides target to cover each designed crRNA (colored rectangles in **Figure 3a, Table S3**). We cross-react the crRNAs with target samples and no target samples in a total of 30 reactions to validate the assay’s specificity. **Figure 3b** shows the fluorescent intensity over 60 minutes of Cas13a reactions. An increase in fluorescent intensity was only observed in cases where targets and crRNAs were matched, confirming the assay’s specificity. In the case of crRNA 3, no significant fluorescent signal increase was observed, which is likely due to the low trans-cleavage activity ^33^. In addition, crRNA1 and 4 showed the highest trans-cleavage activity among the cases where the higher fluorescent intensity was observed after 60 minutes of reaction.

**Figure 3.**
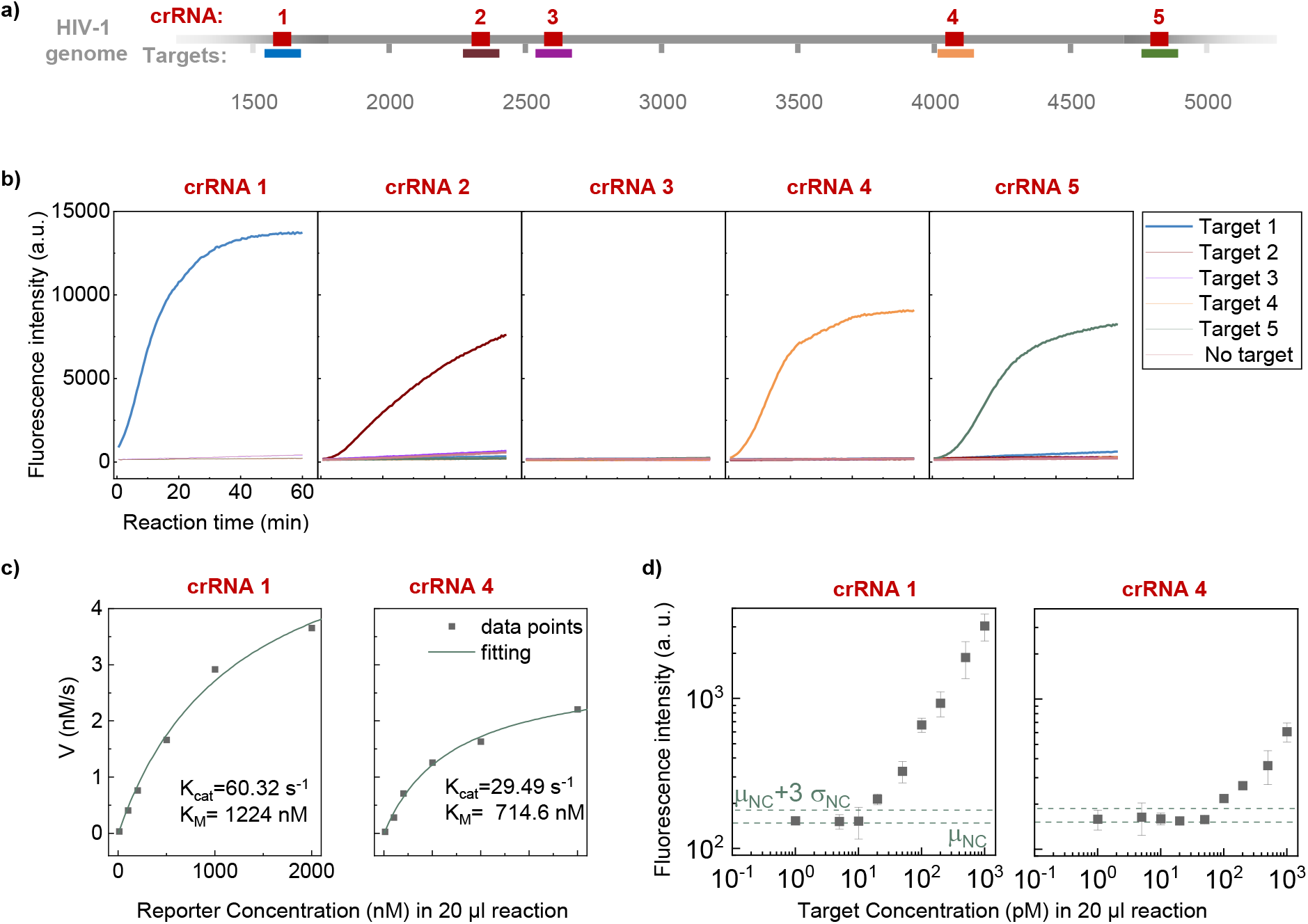
Optimization of Cas13 crRNA and bulk assay characterization. a) Schematic of the HIV-1 genome and the location of each crRNA spacer and the target region. b) Fluorescence intensity values over 60 minutes for 5 different crRNA and their corresponding targets (positive), no-target (NTC), and negative control (NC) samples. c) Michaelis-Menten kinetic study of the Cas13a assay using crRNA 1 and crRNA 4. d) Sensitivity test of CRISPR assay using crRNA 1 and crRNA 4. In each case, three NTC cases were tested to determine the background fluorescent intensity as μ_NTC_+3σ_NTC_, where μ_NTC_ and σ_NTC_ are the averages and standard deviation of the NTC cases, respectively.

To further compare the performance of the Cas13a assay using crRNA 1 and 4, we performed a Michaelis-Menten kinetic study on the system. **Figure 3c** presents the measurements of reaction rates for the trans-cleavage activity of Cas13a proteins for crRNA1 and 4. Each data point is a measured initial reaction velocity (nM/s) for a titrated reporter concentration. **Figure S4** shows the details of cleaved reporter concentration and measurements of cleavage speed. In order to extract the kinetic properties of Cas13 proteins using crRNA1 and 4, the curves in **Figure 3d** were fitted using nonlinear regression based on the Michaelis-Menten equation:

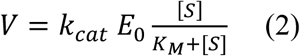

where *E*_0_ is the target-activated Cas/13−cRNA complex concentration, [*S*] is the reporter concentration, *k*_cat_ is the catalytic turnover rate of the enzyme, and *K*_M_ is Michaelis constant. For the reaction using crRNA 1 and 4, the catalytic rate of 29.49 s^-1^ and 60.32 s^-1^ were measured, respectively. The assay using crRNA1 displayed a reaction with a higher cleavage rate. Therefore, crRNA1 was chosen for our digital assay to obtain a faster signal. In addition, we also quantified the bulk assay limit of detection using crRNA 1 and 4. As shown in **Figure 3d**, HIV-1 Cas 13 assay using crRNA1 showed a better limit of detection of ∼20 pM compared to ∼100 pM when using crRNA4. The pM range of limit of detection in an amplification-free bulk Cas13 assay is on par with the previously reported Sherlock assay (∼50 pM) ^15^.

### Optimizing HIV-1 STAMP-dCRISPR assay time

To obtain the optimal reaction time for HIV-1 assay, we measured the *PPR* at different reaction times for the Cas13a assay containing 5 fM HIV-1 synthetic RNA. **Figure 4a** presents the fluorescence images at various reaction times. As the reaction time increased, more positive pores were observed in fluorescent images. This happened because more reporters would be degraded in the positive pores as the reaction time increases, resulting in more wells reaching fluorescent intensity above the sensor detection sensitivity. **Figures 4b** and **Figures 4c** show the corresponding fluorescent intensity (FI) of positive and negative pores and their distributions, respectively. These results confirmed our observation that more positive pores were detected as the reaction time increased. To quantify the effect of reaction time, *PPR* was plotted from 0 to 60 minutes of reaction (**Figures 4d** and **Table S4**). As expected, *the PPR* increases as time passes; however, the ratio becomes stable after 30 minutes. This means that the shortest time to develop a reliable *PPR* reading is about 30 minutes in our assay.

**Figure 4.**
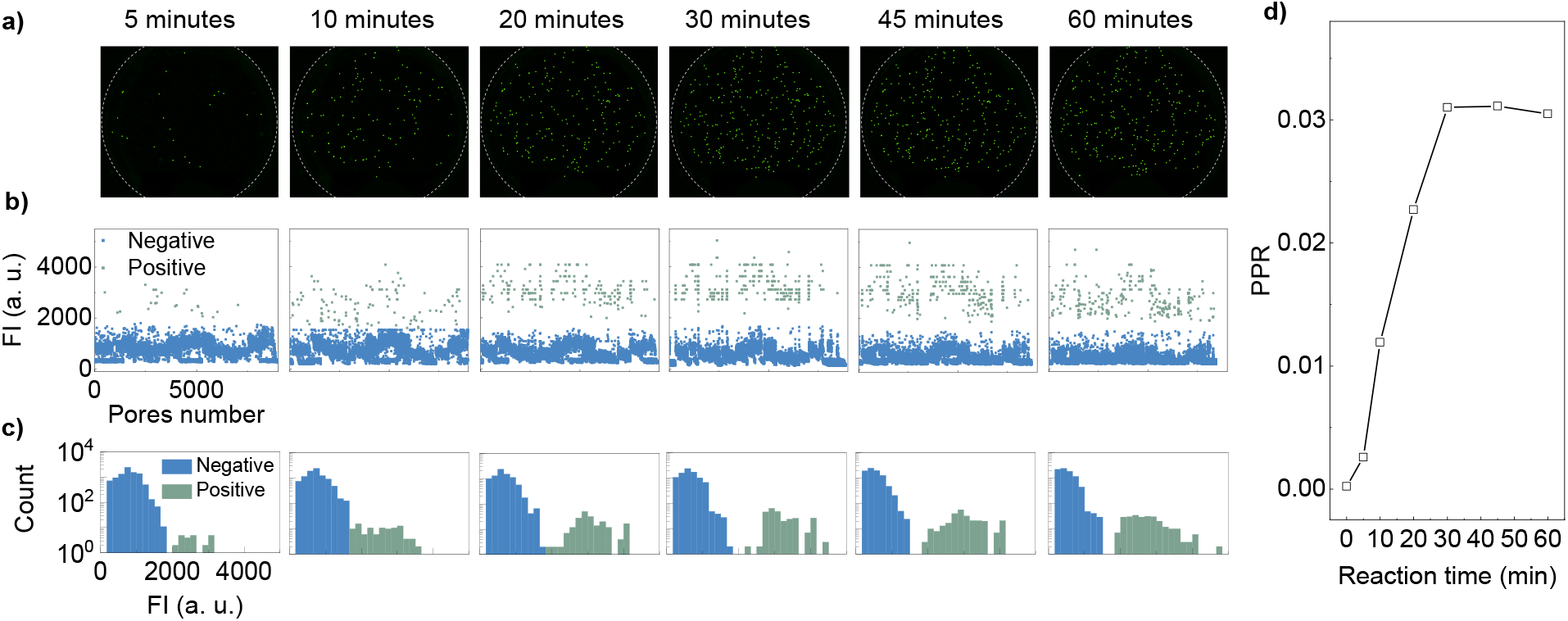
Optimizing STAMP-dCRISPR assay time. a) Fluorescent images illustrating the positive and negative pores at different reaction times from 0 to 60 minutes. The dashed grey circles illustrate the membrane edge. All positive pores are labeled with a filled green circle for better demonstration. b) Fluorescent intensity inside all filled pores (positive and negative) at different reaction times. Positive and negative pores are labeled as green and blue circles, respectively. c) Distribution of fluorescent intensity emitted from positive (green bars) and negative (blue bars) pores. d) The ratio of positive pores (*PPR*) at different reaction times.

Based on the measured *k*_cat_ (**Figure 3c)**, a single activated Cas13a enzyme would produce ∼13 nM of cleaved reporters (fluorescent probes) inside each pore (volume of 13 pL) in a 30 minute reaction. In contrast, a bulk reaction of 20 μL volume would only have produced ∼ 9 pM with the same 30 min reaction. Decreasing the reaction volume from microliter to picolitre would increase the fluorescent concentration by around 6 orders of magnitude and thus help improve the lower limit of detection.

### Analytical performance test with synthetic HIV-1 RNAs

A series of synthetic HIV-1 RNA dilutions from 10 aM to 5 pM were tested to examine the quantitative analytical performance of the STAMP-dCRISPR. **Figure 5a** presents the fluorescent images at different HIV-1 RNA concentrations. As expected, more positive pores were detected as HIV-1 RNA concentration increased. The *PPR* at different target concentrations is plotted in **Figure 5b**. Expectedly, the measured *PPR* increased from 3.7×10^−4^ at 10 aM to 0.99 at 5 pM. These results revealed the dynamic range of STAMP-dCRISPR from 100 aM to 1 pM (4 orders of magnitude). For a concentration lower than 100 aM, the background noise of the system would interfere with the quantification, and accurate measurement could not be achieved. On the other hand, the *PPR* would saturate at 1 for cases with a concentration higher than 1 pM.

**Figure 5.**
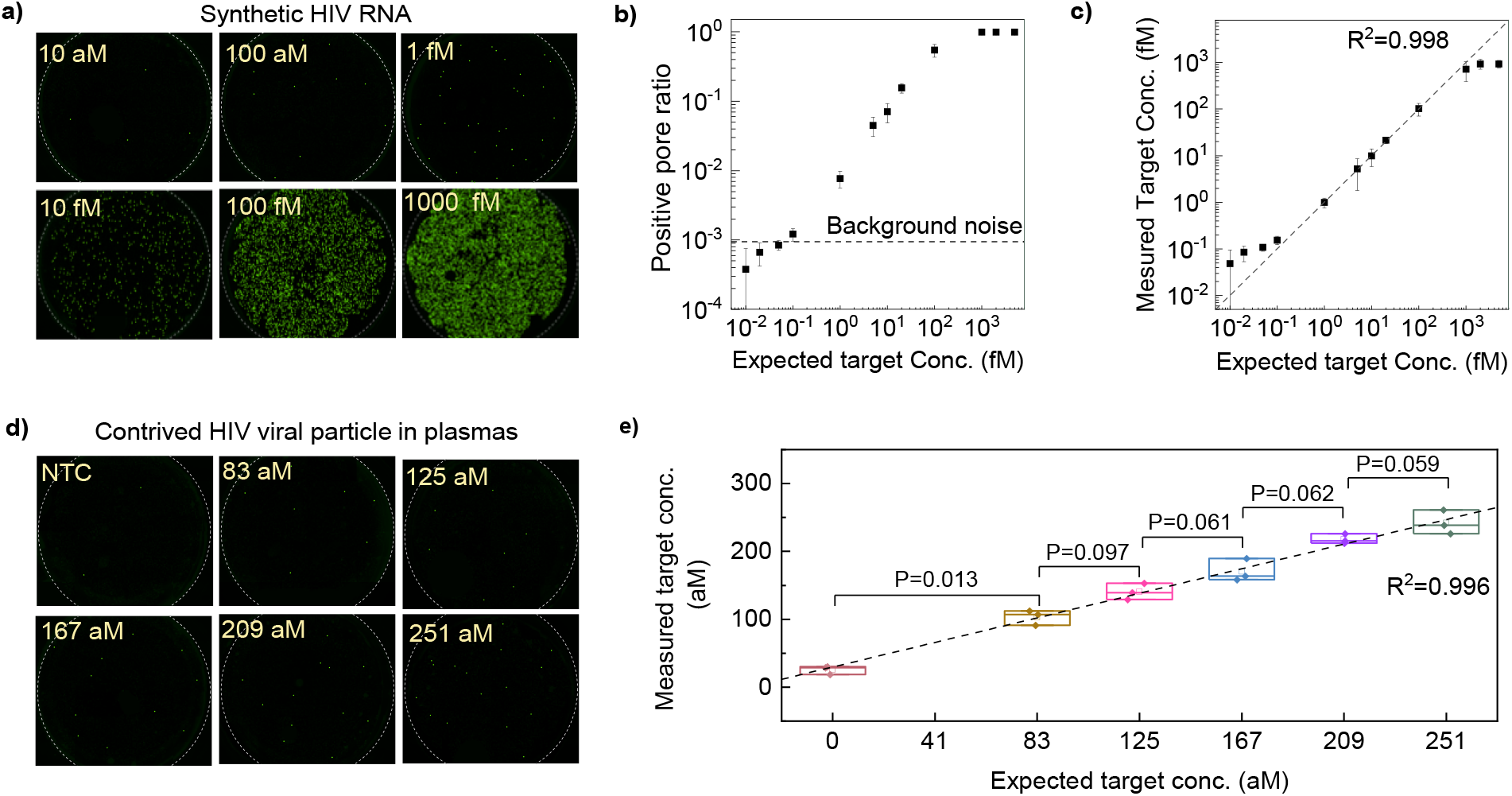
Analytical performance test using synthetic samples and resolution test using contrived plasma sample. a) Representative fluorescent images illustrating positive and negative pores at different target concentrations from 10 aM to 1 pM. All positive pores are labeled with a filled green circle for better demonstration. b) The measured positive pore ratios (PPR) at different HIV-1 RNA concentrations from 10 aM to 5 PM. The top dashed line represents the background noise. c) Comparison of measured HIV RNA concentrations to the expected concentrations. d) Representative fluorescent images illustrating positive and negative pores at different HIV spiked samples. All positive pores are labeled with a filled green circle for better demonstration. e) Measured target concentration of the spiked samples along with the *p* values obtained from a t-test between adjacent cases.

**Figure 5c** presents the measured concentrations via STAMP-dCRISPR versus the expected target concentration (**Table S5**). The measured concentrations in the dynamic range (100 aM to 1 pM) agree very well with the expected concentrations (R^2^=0.998), confirming the absolute quantification capability of the STAMP-dCRIPSR. With the background noise defined as μ_NTC_+3σ_NTC_, the LOD of the STAMP-dCRIPSR was determined to be around 100 aM. As compared to the LOD of 20 pM in the bulk assay shown in **Figure 3d**, the STAMP-dCRIPSR improved the LOD by 5 orders of magnitude.

### Resolution test with contrived plasma sample

To examine the resolution of viral load from the plasm samples, we prepared a serial of mock plasma samples with HIV-1 viral particles spiked into the healthy plasma. Each concentration was prepared in triplicates. The viral RNAs were extracted from these mock plasma samples using a column-based extraction process before being quantified using STAMP-dCRISPR. **Figure 5d** presents six representative fluorescent images. It is evident that more positive pores were observed as we increased the viral concentration. **Figure 5e** shows the measured concentration obtained by STAMP-dCRISPR versus the expected values (see **Table S5** for testing statistics). The measured concentrations agree very well with the input concentrations (R^2^=0.996), confirming the capability of STAMP-dCRISPR system for the absolute quantification of plasma samples. In addition, the *p*-value obtained from the t-test between adjacent concentration revealed that the STAMP-dCRISPR could differentiate the spiked samples with a resolution of 42 aM with >90% confidence. This resolution is equivalent to resolving 5 copies of HIV-1 RNAs in the STAMP device.

### Clinical validation of STAMP-dCRISPR

To demonstrate the clinical utility of the STAMP-dCRISPR, we tested 20 clinical HIV plasma samples using STAMP-dCRISPR. Like the mock samples, we extracted the HIV viral RNA using a column-based extraction process. The extracted RNA was aliquoted into two identical duplicates and tested with STAMP-dCRISPR and RT-PCR, respectively. **Figure 6a** shows the fluorescent images of STAMP-dCRISPR results. **Figure 6b** presents the real time RT-PCR results with six concentration references (R1-R6). The *C*_t_ values showed a linear logarithmic relationship with the reference concentrations, which validates the RT-PCR assay (**Figure S5**).

**Figure 6.**
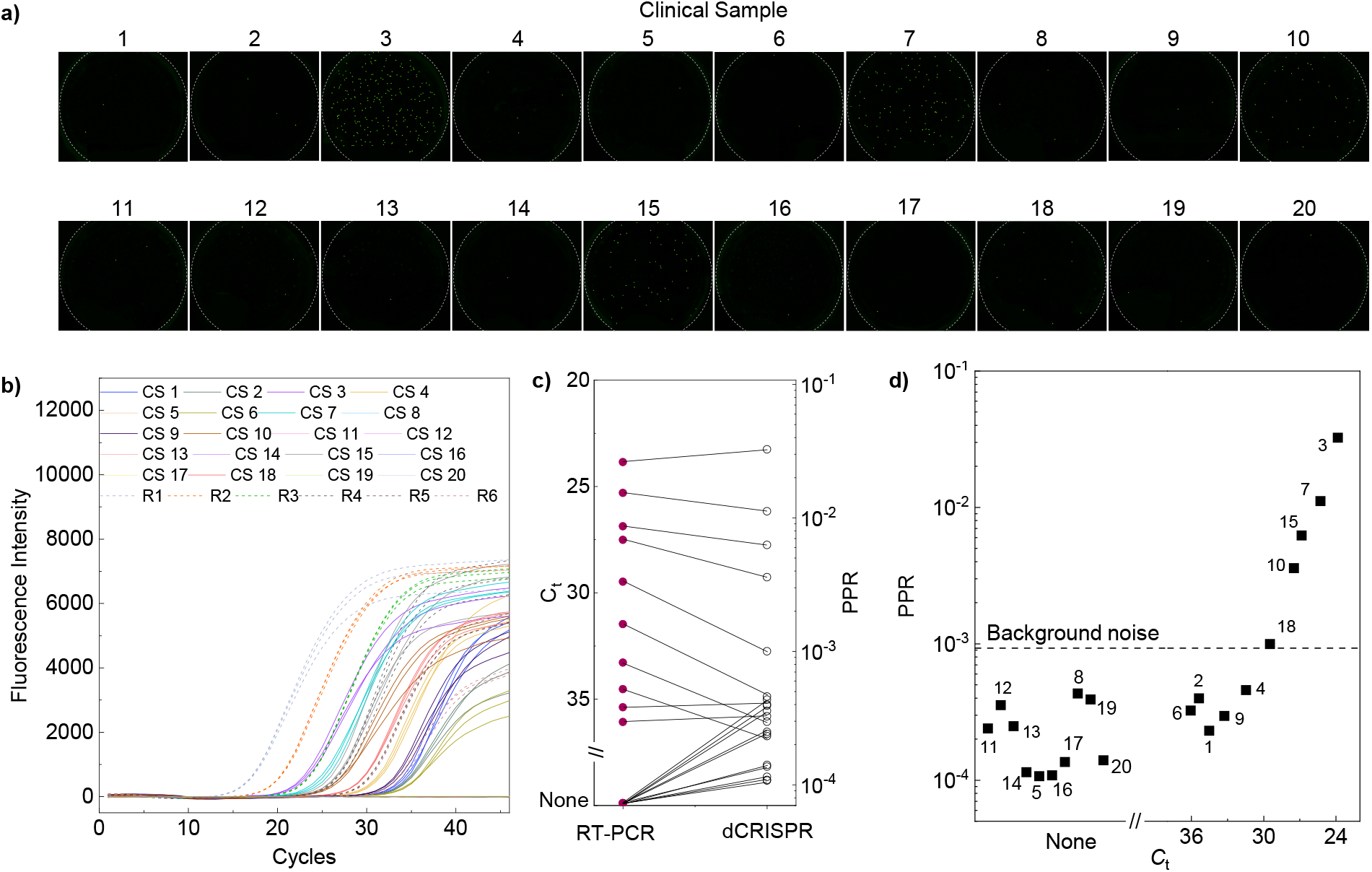
Clinical samples test using STAMP-dCRISPR. a) Fluorescent images illustrating positive and negative pores for different clinical samples. All positive pores are labeled with a filled green circle for better demonstration. b) Fluorescent intensity of 20 clinical samples and 6 analytical samples during 46 cycles of RT-PCR amplification. c) Correspondence between the *C*_t_ values obtained from RT-PCR and PPR values from STAMP-dCRISPR. d) *C*_t_ values versus PPR values for all 20 clinical samples. *C*_t_ values for non-amplified cases were set as None in the figure.

To benchmark the STAMP-dCRISPR results with the RT-PCR, *C*_t_ values of RT-PCR results were compared with the PPR values measured by STAMP-dCRISPR (**Figure 6c** and **Figure 6d**). The *PPR* corresponded well with *C*_t_ values for RT-PCR positive cases and showed the potential of STAMP-dCRISPR for clinical HIV detection. In addition, The *PPR* was less than background noise in all 10 RT-PCR negative cases (no *C*_t_ values), and no false positive results were observed using STAMP-dCRISPR.

## Discussion

In just 5 years, the CRISPR diagnostics has expanded, growing from a set of molecular biology discoveries to multiple FDA-authorized COVID-19 tests ^11^. Digital CRISPR diagnostics provide high standards for next-generation CRISPR diagnostic platforms, such as amplification and calibration-free quantification and single-nucleotide specificity ^38,45^. Here, we developed a membrane-based digital CRISPR-Cas13a system to not only quantify viral loads of HIV-1 but also provide a simple and inexpensive platform for digital CRISPR diagnostics. Unlike previous digital CRISPR systems, our platform does not require complex digitalization systems such as droplet emulsion ^34,45^ techniques or micro and nanofabrication ^38^. A commercial PCTE membrane that is widely available and inexpensive would be a powerful tool for pathogen detection and quantification. In addition, our simple and novel stamping technique will provide a great platform for users without expertise in microfluidics to work with these membranes.

Two crucial factors could be further optimized to shorten the CRISPR assay turnaround time. One is the trans-cleavage speed of Cas proteins and the other is the highly sensitive sensors to read out the signals. Multiple research studies have shown that the crRNA sequence ^32,33^ and engineered Cas13a proteins itself ^64^ could drastically affect the trans-cleavage speed of Cas proteins. We observed a similar behavior where the fluorescent signal produced by the Cas13a proteins could be improved by 2 orders of magnitude by optimizing the crRNA sequences.

In STAMP-dCRISPR, a membrane that contains ∼ 1.32×10^4^ pores with an average volume of 13 pL was utilized. Based on the Poisson statistics (Eq.1), the minimum and maximum detectable concentrations using STAMP-dCRISPR would be 9.58 aM and 1.21 pM, respectively. Based on our experimental results, STAMP-dCRISPR showed a dynamic range between 100 aM and 1 pM. While our upper limit of the obtained dynamic range was close to theory, the lower limit was affected by the background noise. Engineering the pore volume and number of pores could further improve the LOD and the dynamic range of STAMP-dCRISPR.

Finally, absolute quantification of HIV clinical samples was carried out to evaluate STAMP-dCRISPR performance. We observed 100% specificity compared with the RT-PCR result, and no false positive result was detected. However, the 5 PCR positive samples with *C*_*t*_ values lower than 30 were not picked up by STAMP-dCRISPR (**Figure 6d)**. This is mainly due to the subsampling error, which arise in all biological assays (digital or analog) which does not analyze the full volume of sample ^65^. While the CRISPR mixture volume is 20 μL, the STAMP device takes ∼ 100 nL for each analysis. Subsampling error could affect the lower detection limit at low concentrations and is independent of the instrumentation and signal acquisition. Several methods could alleviate this challenge in the future. Using multiple membranes for a single sample could increase the volume of testing samples and substantially improve the system performance. Also, magnetic-bead based preconcentration ^47^ could be utilized to enrich target RNA molecules inside the membrane.

## Methods

### STAMP device fabrication

The PMMA holders were prepared by cutting the PMMA sheets with 1/8” thickness using a laser cutter machine (Universal Laser System). Two pieces of PMMA with the dimensions of 24×24 mm and 35×35 mm with inner circles of 11 and 13 mm were fabricated and attached using acrylic cement (United States Plastic Corporation, cat# 46872). To handle the track-etched polycarbonate membranes (Sterlitech Corporation, cat# PCT25025100), we utilized a vacuum pen (Pen-vac pro series V8910). The PVP layer of the membranes was removed by dipping the membranes in 10% acetic acid for 30 min, followed by heating to 140 °C for 60 min in a vacuum oven. Afterward, the membrane was attached to the holder using adhesive tapes (70 μm thickness). We used mineral oil purchased from Sigma-Aldrich (cat# 69794-500ML) to seal the membrane.

### Automated data acquisition and analysis

The fluorescent images were taken using a fluorescent microscope (Nikon ECLIPSE Ti). The integration time was set as 6 s to image the membrane. To cover the whole membrane, a motorized stage (Prior OptiScan) was utilized, and 24 images were taken to cover the membrane. A stitching algorithm was employed to obtain the whole fluorescent images of each membrane. Afterward, MATLAB (MathWorks) software was used to implement a *k*-means clustering algorithm to differentiate between positive and negative pores (See **Figure S3** for details of stitching and clustering algorithm).

### crRNA design and selection

The optimal protospacer length observed for Cas13a is 28 nucleotides along ^66^. In addition, Abudayyeh et al. analyzed the flanking regions of protospacers and found that sequences starting with a G immediately after the 3’ end of the protospacer were less effective relative to all other nucleotides (A, U, or C) ^67^. Therefore, considering the protospacer-flanking site (PFS), 28 nucleotide crRNA protospacer sequences were designed by targeting the HIV-1 type B sequence downloaded from the NCBI website. In the next step, 496 complete HIV sequences deposited in NCBI server were downloaded on 9/14/2021. These sequences were aligned using SnapGene software. We searched designed crRNAs against the aligned sequence with more than 80% similarity and chose five matched crRNAs (**Table S1**). It should be mentioned that we used a previously validated sequence for the direct repeat region of the crRNA as follows: 5’-GAUUUAGACUACCCCAAAAACGAAGGGGACUAAAAC -3’ ^23^.

### Cas13 reaction mixture

The designed crRNAs were synthesized by Integrated DNA Technologies. The crRNAs were resuspended in pH 7.5 buffer and stored at -80 °C. LwaCas13a proteins were purchased from MCLAB (cat# CAS13a-100). Cas13a and crRNA were mixed in 1×PBS to form the non-activated Cas13a/crRNA at room temperature for 20 min and stored at -80°C. In the cleavage reaction, the 2 μL of non-activated Cas13a/crRNA complex was mixed with the 2 μL of RNA target in 9.5 μL of water, 0.5 of Murine RNase Inhibitor (purchased from New England Biolabs, cat# M0314S), 2 μL of FQ-labeled reporter (RNaseAlert substrate purchased from IDT, cat# 11-04-02-03), and 4 μL of a CRISPR buffer consisting of 20 mM HEPES-Na pH 6.8, 50 mM KCl, 5 mM MgCl_2_, and 5% glycerol. Afterward, the mixed solution was incubated in a microplate (for bulk assays) or PCTE membranes (for digital assays) at 37°C for different reaction times.

### Bulk Cas 13a assay

For bulk assay fluorescent signal acquisition, the 20 μL CRISPR reaction mix was incubated inside a 384 well black plate (purchased from ThermoFisher, cat# 142761). The fluorescent signal was measured every 30 s for different reaction times using a microplate reader (Tecan plate reader infinite 200 PRO). The excitation wavelength was set as 480 nm with a bandwidth of 9 nm, and the emission wavelength was set as 530 nm with a bandwidth of 20 nm. The temperature was fixed at 37°C. The gain and integration time were set as 110 and 20 μs, respectively.

### STAMP-dCRISPR assay

For the STAMP-dCRISPR, the CRISPR reaction mix was incubated inside the PCTE membrane using STAMP. In the first step, 8 μL of reaction mix was dropped on top of a glass surface, and the stamp was slowly placed on top of it. After 60 seconds, we started the second step and sealed the top surface of the membrane by adding 60 μL of mineral oil on top of the membrane. The stamp was peeled off from the glass surface in the third step. In the final step, the stamp was placed on top of the system base, which consisted of a glass substrate and a double-sided tape filled with mineral oil to seal the bottom side of the membrane. The sealed system was placed on top of a hot plate (Fisherbrand Isotemp Hot Plate) at 37 °C for different reaction times.

### Contrived plasma mock sample and clinical samples

To prepare the contrived plasma mock sample, different copies (1000, 1500, 2000, 2500, and 3000) of HIV viral particles (Seraseq, cat# 0740-0004) were mixed in 140 μL of fresh, healthy plasma (Research Blood Components). After mixing, the samples were preserved at -80 °C. The clinical HIV plasma samples were obtained from Hershey Medical Center by an approved institutional review board (IRB) of the Pennsylvania State University. All samples were coded to remove information associated with patient identifiers. The plasma samples were stored at -80 °C before the examination.

### Viral RNA extraction from plasma

To extract the RNA a column-based RNA extraction kit purchased from Qiagen (cat# 52904) was utilized. The procedure is optimized for samples with a volume of 140 μL. The sample is first lysed under the highly denaturing conditions provided by a viral lysis buffer. In addition, we added carrier RNA to the lysis buffer, which enhances the binding of viral RNA to the kit membrane and reduces the chance of viral RNA degradation. Afterward, the purification was carried out in 3 steps using a standard centrifuge (Eppendorf centrifuge 5425). We washed the sample using ethanol and 2 washing buffers provided by the kit. In the final stage, we used 10 uL of nuclease-free water (BioLabs, cat# 52904B1500S) as an elution buffer to obtain the extracted RNAs from the membrane.

### HIV-1 RT-PCR assay

We used a one-step, two-enzyme RT-PCR protocol for HIV-1 assays. The reaction has a total volume of 20 μL, consisting of 5 μL TaqMan Fast Virus 1-Step Master Mix (cat# 4444432, Thermofisher), 1.2 μL forward primer (0.6 μm), 1.2 μL reverse primer (0.6 μm), 0.5 μL probe (0.25 μm), and 10 μL RNA templates as well as 2.1 μL PCR grade water. We used a previously validated HIV-1 RT-PCR primer set (Forward primer: 5’-CATGTTTTCAGCATTATCAGAAGGA -3’, and Reverse primer: 5’-TGCTTGATGTCCCCCCACT -3’) ^68^. In addition, the probe was selected as 5’-FAM-CCACCCCACAAGATTTAAACACCATGCTAA-Q -3’, where Q indicates a 6-carboxytetramethylrhodamine group quencher conjugated through a linker arm nucleotide. The following thermal cycling sequences performed the RT-PCR: 50 °C for the first five minutes without repeating to reverse transcription reactions which convert HIV-1 RNA into cDNA, then 95? for 20 seconds without repeating to initiate amplification, followed by 46 cycles of amplification stage consisting of 3 seconds of 95 °C and 30 seconds of 60°C thermal-cycling.

## Data availability

All source data used for generating graphs and charts in main and supplementary figures are included in Supplementary Data. The reference genomes of HIV are downloaded from the NCBI genome database (https://www.ncbi.nlm.nih.gov/genbank). The raw and analyzed datasets generated during the study are available for research purpose from the corresponding author on reasonable request.

## Acknowledgments

This work was partially supported by the National Institutes of Health (R61AI147419) and National Science Foundation (1902503, 1912410, 2045169). Any opinions, findings, and conclusions or recommendations expressed in this work are those of the authors and do not necessarily reflect the views of the National Science Foundation and National Institutes of Health.

## Author contributions

W.G. conceived the concept and supervised the study. R.N., X.L.L. and Y.J. designed the Cas13a assay. R.N, A.J.P, and Y.J. validated the Cas13a assay. R.N. performed the digital assay and analyzed the data. R.N. and T.L. designed and performed the RT-PCR. J. N., W. Greene collected and tested the clinical samples. W.G. and R.N. co-wrote the manuscript, with discussion from all authors.

