## Supplementary information for "STAMP-Based Digital CRISPR-Cas13a (STAMP-dCRISPR) for Amplification-Free Quantification of HIV-1 Plasma Viral Load"

This PDF file includes:

Figures S1 to S5

Tables S1 to S8

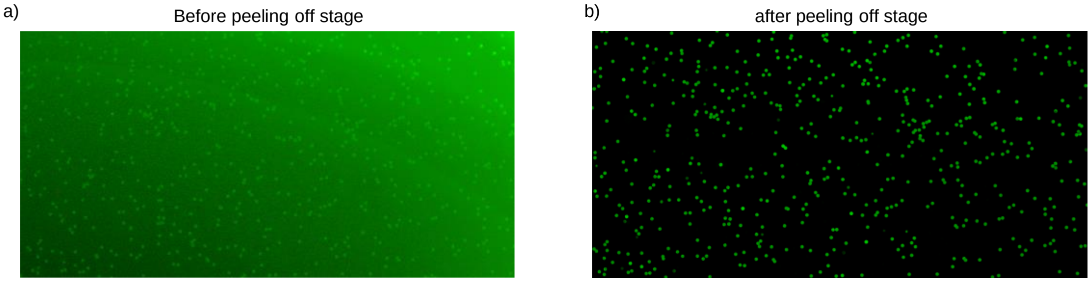


**Figure S1**. Effect of peeling-off stage on the excess sample removal process. Fluorescent images of the membrane a) before and b) after the peeling-off stage of the stamping process. These images confirmed the excess sample removal process in the peeling-off stage.

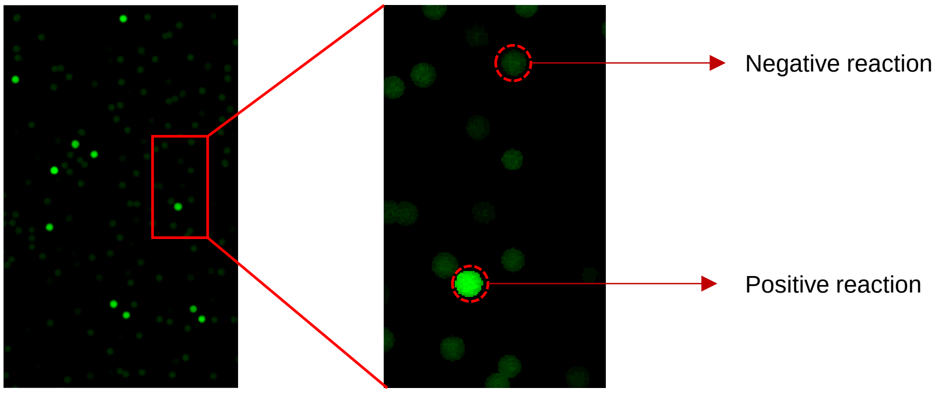


**Figure S2.** Typical fluorescent signals from a negative and positive reaction. Fluorescent image of part of a membrane demonstrating typical fluorescent signals from a negative and positive reaction. The negative reaction containing the unquenched FAM reporters also shows a weak fluorescence signal in our fluorescent images.

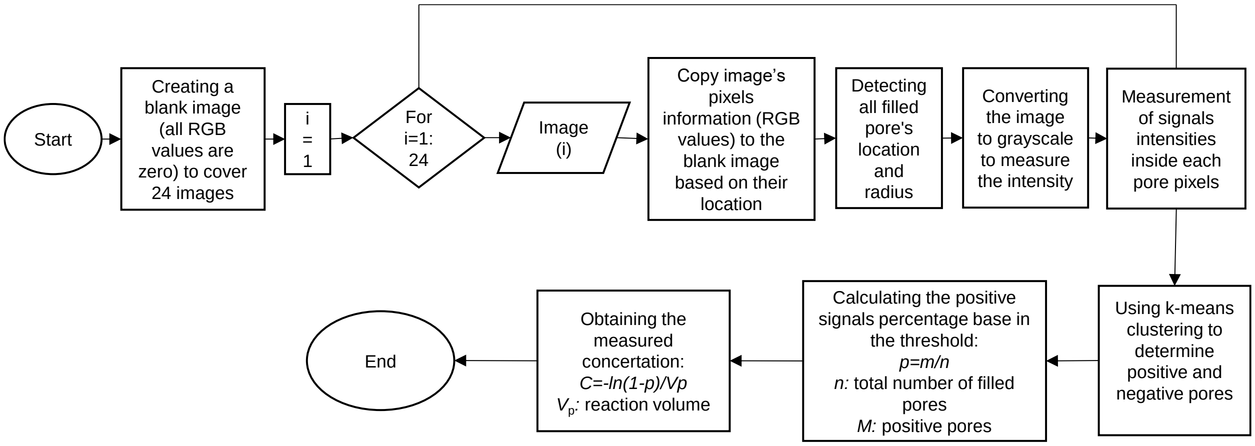


**Figure S3.** Algorithm for acquiring the fluorescent intensity and stitching of the images. The algorithm runs through 24 images and identifies the pixels inside the filled pores in each case. Afterward, The fluorescent intensity of each pore is calculated by combining the intensity of the pixels inside the pores. After examining all 24 images, a k-means algorithm would classify the positive and negative pores based on the fluorescent intensity. Finally, the concentration of the sample would be calculated using Poisson statistics.

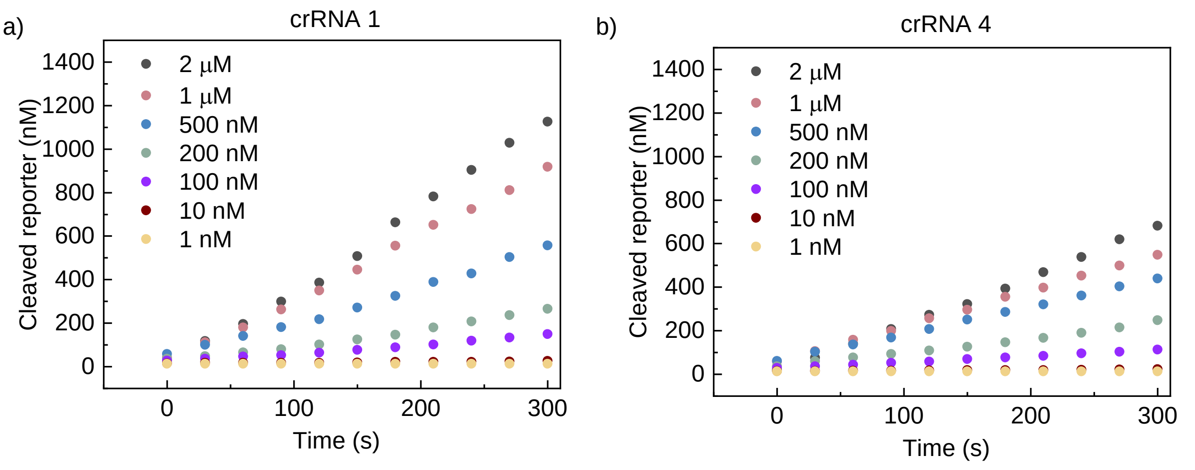


c)

| Reporter concentration (nM) | Cleavage Velocity using crRNA 1 (nM/s) | Cleavage Velocity using crRNA 4 (nM/s) |
|---|---|---|
| 2000 | 3.657189 | 2.208361 |
| 1000 | 2.917791 | 1.634262 |
| 500 | 1.664232 | 1.261519 |
| 200 | 0.762344 | 0.709897 |
| 100 | 0.410205 | 0.283772 |
| 10 | 0.035589 | 0.028096 |

**Figure S4.** Cleavage velocity measurements using crRNA1 and 4. Measurements of cleaved reporters associated with the trans-cleavage activity of the CRISPR Cas13 proteins using a) crRNA 1 and b) crRNA 4. In each case, cleaved reporters were measured for 420 s to extract the cleavage speed (*V* (nM/s)). c) Details of measured cleavage speed of CRISPR Cas13 proteins using crRNA 1 and crRNA 4 at different reporter concentrations.

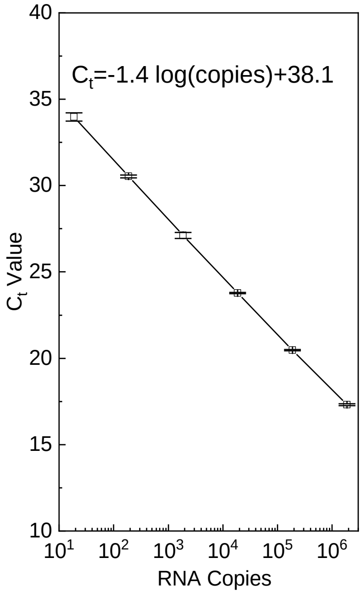


**Figure S5**. The $C_t$ values obtained from the RT-PCR versus the target RNA copies. $C_t$ values showed a logarithmic relationship with the sample concentration and verified our assay.

**Table S1.** The total number of filled and positive pores and the measured positive pore ratio for 4 negative control cases. The average and standard deviation of the positive pore ratio in negative control cases were measured as 0.00033 and 0.0002, respectively. Therefore, our threshold for negative samples was set as $\mu_{NTC}+3\sigma_{NTC}=0.00093$.

| Case Id | Total filled number of wells | Number of positive wells | PPR |
|---|---|---|---|
| 1 | 8652 | 2 | 0.000231163 |
| 2 | 9321 | 3 | 0.000321854 |
| 3 | 8923 | 1 | 0.000112072 |
| 4 | 9145 | 6 | 0.000656096 |

**Table S2.** Detailed sequences of crRNAs

| Description | Sequence |
|---|---|
| HIV-1 crRNA 1 | GAUUUAGACUACCCCAAAAACGAAGGGGACUAAAACGGCUAUACAUUCUUACUAUUUUAUUUAA |
| HIV-1 crRNA 2 | GAUUUAGACUACCCCAAAAACGAAGGGGACUAAAACAAACCUCCAAUUCCCCCUAUCAUUUUUG |
| HIV-1 crRNA 3 | GAUUUAGACUACCCCAAAAACGAAGGGGACUAAAACGCUUUUAUUUUUUCUUCUGUCAAUGGCC |
| HIV-1 crRNA 4 | GAUUUAGACUACCCCAAAAACGAAGGGGACUAAAACCUAAUUUAUCUACUUGTTCAUUUCCUCC |
| HIV-1 crRNA 5 | GAUUUAGACUACCCCAAAAACGAAGGGGACUAAAACUCCCUGUAAUAAACCCGAAAAUUUUGAA |

**Table S3.** Detailed sequences of targets

| | |
|---|---|
| HIV-1 Target 1 fragment | TAGGAGAAATTTATAAAAGATGGATAATCCTGGGATTAAATAAAATAGTAAGAATGTATAGCCCTACCAGCATTCTGGACATAAGACAAGGACCAAAG |
| HIV-1 Target 2 fragment | ATTAGAAGAAATGAGTTTGCCAGGAAGATGGAAACCAAAAATGATAGGGGGAATTGGAGGTTTTATCAAAGTAAGACAGTATGATCAGATACTCATAG |
| HIV-1 Target 3 fragment | AAAGCCAGGAATGGATGGCCCAAAAGTTAAACAATGGCCATTGACAGAAGAAAAAATAAAAGCATTAGTAGAAATTTGTACAGAGATGGAAAAGGAAG |
| HIV-1 Target 4 fragment | TCTATCTGGCATGGGTACCAGCACACAAAGGAATTGGAGGAAATGAACAAGTAGATAAATTAGTCAGTGCTGGAATCAGGAAAGTACTATTTTTAGAT |
| HIV-1 Target 5 fragment | ACAAACTAAAGAATTACAAAAACAAATTACAAAAATTCAAAATTTTCGGGTTTATTACAGGGACAGCAGAAATCCACTTTGGAAAGGACCAGCAAAGC |

**Table S4.** Details information for the total number of filled pores, positive pores, and positive pore ratio at different reaction times.

| Reaction time (min) | Total filled number of filled pores | Number of positive pores | Positive pore ratio (PPR) |
|---|---|---|---|
| 0 | 2 | 9421 | 0.000212292 |
| 5 | 24 | 9235 | 0.002598809 |
| 10 | 110 | 9230 | 0.01191766 |
| 20 | 202 | 8901 | 0.022694079 |
| 30 | 275 | 8872 | 0.030996393 |
| 45 | 271 | 8710 | 0.031113662 |
| 60 | 257 | 8431 | 0.030482742 |

**Table S5.** Details information for the total number of filled pores, positive pores, positive pore ratio, the expected number of targets in each pore (λ), and measured concentrations at different expected target concentrations.

| Expected Target Concentration (fM) | Total number of filled pores | Number of positive pores | Positive pore ratio | Measured Concentration (fM) |
|---|---|---|---|---|
| First batch | | | | |
| 0.01 | 8625 | 4 | 0.000463768 | 0.06 |
| 0.02 | 7835 | 4 | 0.00051053 | 0.07 |
| 0.05 | 7545 | 7 | 0.000927767 | 0.12 |
| 0.1 | 9231 | 10 | 0.001083306 | 0.14 |
| 1 | 8729 | 57 | 0.006529958 | 0.86 |
| 5 | 7342 | 280 | 0.038136747 | 2.99 |
| 10 | 6686 | 400 | 0.059826503 | 7.88 |
| 20 | 5451 | 912 | 0.167308751 | 23.32 |
| 100 | 6925 | 4136 | 0.597256318 | 116.17 |
| 1000 | 8765 | 8722 | 0.995094124 | 797.74 |
| 2000 | 6950 | 6941 | 0.998705036 | 849.34 |
| 5000 | 8574 | 8571 | 0.999650105 | 1016.34 |
| Second batch | | | | |
| 0.01 | 9053 | 5 | 0.000552303 | 0.07 |
| 0.02 | 7432 | 6 | 0.00080732 | 0.10 |
| 0.05 | 8312 | 7 | 0.000842156 | 0.11 |
| 0.1 | 8653 | 12 | 0.001386802 | 0.18 |
| 1 | 8981 | 81 | 0.00901904 | 1.15 |
| 5 | 9232 | 503 | 0.054484402 | 7.13 |
| 10 | 9367 | 766 | 0.081776449 | 11.96 |
| 20 | 8869 | 1271 | 0.143308152 | 19.75 |
| 100 | 8882 | 4150 | 0.467237109 | 80.43 |
| 1000 | 9080 | 9069 | 0.998788546 | 857.85 |
| 2000 | 9456 | 9453 | 0.999682741 | 1029.17 |
| 5000 | 8938 | 8931 | 0.999216827 | 956.21 |
| Third batch | | | | |
| 0.01 | 8322 | 1 | 0.000120163 | 0.016 |
| 0.02 | 7325 | 5 | 0.000682594 | 0.09 |
| 0.05 | 7880 | 6 | 0.000761421 | 0.10 |
| 0.1 | 9321 | 11 | 0.001180131 | 0.15 |
| 1 | 10755 | 80 | 0.007438401 | 0.95 |
| 5 | 9242 | 390 | 0.042198658 | 5.53 |
| 10 | 7329 | 623 | 0.085004776 | 11.34 |
| 20 | 6408 | 826 | 0.128901373 | 17.62 |
| 100 | 8713 | 5007 | 0.574658556 | 109.32 |
| 1000 | 8425 | 8235 | 0.977448071 | 484.36 |

| | | | | |
|---|---|---|---|---|
| 2000 | 8133 | 8125 | 0.999016353 | 921.21 |
| 5000 | 7123 | 7111 | 0.998315317 | 815.12 |

**Table S6.** Details information for the total number of filled pores, positive pores, positive pore ratio, and measured concentrations at different input target copies inside the 140 µL of plasma.

| Input targets (Copies) | Total number of filled pores | Number of positive pores | Positive pore ratio | Measured targets (Copies) |
|---|---|---|---|---|
| | | First batch | | |
| 0 | 8845 | 2 | 0.000226116 | 347 |
| 1000 | 8423 | 6 | 0.000712335 | 1096 |
| 1500 | 7510 | 9 | 0.001198402 | 1844 |
| 2000 | 8585 | 11 | 0.001281305 | 1972 |
| 2500 | 8428 | 14 | 0.00166113 | 2557 |
| 3000 | 9041 | 16 | 0.001769716 | 2725 |
| | | Second batch | | |
| 0 | 6752 | 1 | 0.000148104 | 227 |
| 1000 | 9115 | 8 | 0.000877674 | 1350 |
| 1500 | 8252 | 9 | 0.001090645 | 1678 |
| 2000 | 8765 | 13 | 0.001483172 | 2285 |
| 2500 | 7921 | 14 | 0.001767454 | 2721 |
| 3000 | 8321 | 17 | 0.002043024 | 3146 |
| | | Third batch | | |
| 0 | 8501 | 2 | 0.000235266 | 361 |
| 1000 | 8346 | 7 | 0.000838725 | 1290 |
| 1500 | 8895 | 9 | 0.001011804 | 1557 |
| 2000 | 8871 | 11 | 0.001239995 | 1908 |
| 2500 | 6521 | 11 | 0.001686858 | 2597 |
| 3000 | 8124 | 16 | 0.001969473 | 2876 |

**Table S7.** Details information for the total number of filled pores, positive pores, positive pore ratio, the expected number of targets in each pore (λ), and measured concentrations of clinical samples.

| Sample # | Total number of filled pores | Number of positive pores | Positive pore ratio | λ |
|---|---|---|---|---|
| 1 | 8651 | 2 | 0.000231187 | 0.000231214 |
| 2 | 7515 | 3 | 0.000399202 | 0.000399281 |
| 3 | 7234 | 235 | 0.032485485 | 0.032912355 |
| 4 | 8723 | 4 | 0.000458558 | 0.000458663 |
| 5 | 9316 | 1 | 0.000107342 | 0.000107348 |
| 6 | 6153 | 2 | 0.000325045 | 0.000325098 |
| 7 | 8424 | 94 | 0.011158594 | 0.012552231 |
| 8 | 6928 | 3 | 0.000433025 | 0.000433119 |
| 9 | 10127 | 3 | 0.000296238 | 0.000296282 |
| 10 | 10870 | 39 | 0.003587856 | 0.003594308 |
| 11 | 8324 | 2 | 0.000240269 | 0.000240298 |
| 12 | 8412 | 3 | 0.000356633 | 0.000356697 |
| 13 | 8021 | 2 | 0.000249345 | 0.000249377 |
| 14 | 8740 | 1 | 0.000114416 | 0.000114423 |
| 15 | 8801 | 55 | 0.00624929 | 0.006268898 |
| 16 | 9201 | 1 | 0.000108684 | 0.00010869 |
| 17 | 7338 | 1 | 0.000136277 | 0.000136286 |
| 18 | 9021 | 9 | 0.000997672 | 0.000998172 |
| 19 | 7668 | 3 | 0.000391236 | 0.000391313 |
| 20 | 7123 | 1 | 0.00014039 | 0.0001404 |

**Table S8.** Details information for the RT-PCR *Ct* values of clinical samples.

| Sample # | Total number of filled pores | Ct Value (mean) | Ct Value (STD) |
|---|---|---|---|
| 1 | 8651 | 34.53 | 0.22 |
| 2 | 7515 | 35.39 | 0.14 |
| 3 | 7234 | 23.86 | 0.09 |
| 4 | 8723 | 31.48 | 0.23 |
| 5 | 9316 | None | None |
| 6 | 6153 | 36.08 | 0.05 |
| 7 | 8424 | 25.3 | 0.2 |
| 8 | 6928 | None | None |
| 9 | 10127 | 33.28 | 0.18 |
| 10 | 10870 | 27.51 | 0.16 |
| 11 | 8324 | None | None |
| 12 | 8412 | None | None |
| 13 | 8021 | None | None |
| 14 | 8740 | None | None |
| 15 | 8801 | 26.87 | 0.11 |
| 16 | 9201 | None | None |
| 17 | 7338 | None | None |
| 18 | 9021 | 29.48 | 0.11 |
| 19 | 7668 | None | None |
| 20 | 7123 | None | None |
